# The lncRNA *Statera* regulates synaptogenesis by repressing the retrotransposon *copia* in *Drosophila*

**DOI:** 10.64898/2026.09.01.748680

**Authors:** Shuhao Wang, Peter M’Angale, Cong Xiao, Gimena Alegre, Alfred Simkin, Anna Malinkevich, Travis Thomson

**Author notes:** Corresponding author. (T.T.).

## Abstract

Retrotransposons are mobile genetic elements that can impair genome integrity, and their insertions and dysregulation have been implicated in various diseases. However, growing evidence suggests that retrotransposons may play important physiological roles. At the *Drosophila* larval neuromuscular junction (NMJ), the ViSyToR (Viral Synaptic Transfer of RNA) pathway requires the active retrotransposon *copia* as a key negative regulator of synaptogenesis. Here, we identify a long non-coding RNA, *Statera* (*Stae*), as a regulator of *copia*, initially discovered through its physical association with the Copia protein. *Stae* is highly expressed in the central nervous system and body wall muscles of *Drosophila* larvae. CRISPR knockout or RNAi knockdown of *Stae* leads to a general decrease of bouton number at the NMJ, indicating its role in promoting synaptogenesis. Furthermore, *copia* is highly upregulated in *Stae* mutants, suggesting that *Stae* functions to repress *copia*. Mechanistically, *Stae* reduces *copia* RNA stability and limits *copia* DNA copy number. Additional knockdown of *copia* in motor neuron-specific *Stae* RNAi animals rescues the abnormal NMJ morphology. Together, our findings reveal an unexpected role for a long non-coding RNA in promoting synaptogenesis by repressing a retrotransposon.

## INTRODUCTION

Transposable elements (TEs) are mobile genetic elements that constitute substantial portions of eukaryotic genomes (Lander et al., 2001; Waterston et al., 2002). They are traditionally thought to be parasitic or non-functional, yet accumulating evidence has shown that they can drive the evolution of new genes with important cellular functions (Bourque et al., 2018; Hayward & Gilbert, 2022). In addition, recent studies have shown that retrotransposons may play direct physiological roles in embryogenesis and neural development (Percharde et al., 2018; Bodea et al., 2024; Chang et al., 2025). In *Drosophila*, the involvement of TEs in neural development is exemplified by the ViSyToR (Viral Synaptic Transfer of RNA) pathway at the larval neuromuscular junction (NMJ). We previously described how the retrotransposon-like protein dArc1 is transported across the NMJ to promote synaptogenesis (Ashley et al., 2018). Recently, we identified the *Drosophila* retrotransposon *copia* as another key component of the ViSyToR pathway and showed that it inhibits synaptogenesis (M’Angale et al., 2025). Similar to dArc1, Copia RNA and protein are transported from pre- to postsynaptic sites in a virus-like manner. Interestingly, Copia and dArc1 have opposing functions and negatively regulate each other (M’Angale et al., 2025). Unlike *dArc1*, however, *copia* is an extant TE, raising the question of how the nervous system harnesses a potentially parasitic element for developmental control.

Long non-coding RNAs (lncRNAs) are RNAs longer than 200 nt that do not encode proteins and are typically transcribed by RNA polymerase II (Nojima & Proudfoot, 2022). LncRNAs were originally regarded as transcriptional noise but have received increasing attention over the past two decades due to their extensive roles in gene regulation. LncRNAs have been reported to regulate diverse biological processes such as chromatin remodeling, transcription, RNA stability, and translation (Marchese et al., 2017). In mammals, lncRNAs MALAT1 and Gomafu promote synaptogenesis by controlling alternative splicing of downstream synaptic genes (Barry et al., 2014; Bernard et al., 2010). The roles of lncRNAs at the *Drosophila* NMJ remain largely unknown. *hsrω*, a stress-responsive lncRNA, promotes synapse formation at motor neuron terminals and locomotor activity in *Drosophila* (Lo Piccolo & Yamaguchi, 2017). However, the detailed mechanism remains to be elucidated. Recently, the lncRNA *SDRG* was reported to regulate synaptic growth at the *Drosophila* NMJ by antagonizing *frequenin 2*, which encodes a calcium-binding protein (Cui et al., 2026).

The long non-coding RNA *Statera* (*Stae*, originally *CR34335*) is an abundantly expressed RNA in *Drosophila* with unknown biological function (Inagaki et al., 2005; Xue et al., 2014). *Stae* expression is upregulated during aging (Davie et al., 2018) and cocaine exposure (Baker et al., 2021), and its RNA is enriched in extracellular vesicles (EVs) from *Drosophila* cell lines (Lefebvre et al., 2016). Notably, TEs were also found to be upregulated or enriched under these conditions (Silva et al., 2026; Ashley et al., 2018; Lefebvre et al., 2016; Li et al., 2013). LncRNAs have been shown to repress TEs in yeast and human cells, potentially by inducing changes in chromatin (Berretta et al., 2008; Zhang et al., 2022). The correlation between *Stae* RNA and TE expression, together with previously reported roles of lncRNAs in TE control, makes *Stae* a promising candidate for investigating TE regulation in the nervous system.

In an effort to identify RNA interactors of dArc1 and Copia proteins in the ViSyToR pathway, we found that *Stae* co-immunoprecipitates with Copia proteins in *Drosophila* S2 cells and body wall muscles (BWMs). Genetic manipulation of *Stae* using CRISPR knockout (KO) or neuronal RNAi knockdown (KD) led to a decrease of synaptic bouton numbers at the NMJ, suggesting that *Stae* regulates synaptogenesis in *Drosophila*. We further showed that *Stae* inhibits *copia* expression using digital PCR (dPCR) and immunostaining. Interestingly, *copia* DNA copy numbers are increased, and *copia* transcript levels persist following actinomycin D (ActD) treatment in *Stae* KO, suggesting increased *copia* RNA stability and retrotransposition in the absence of *Stae*. Collectively, these data provide new evidence that a lncRNA mediates *Drosophila* synapse formation through retrotransposon regulation.

## RESULTS

### *Stae* is expressed in the CNS and BWMs of *Drosophila* larvae

To identify potential candidates in the ViSyToR pathway, we performed an unbiased screen for RNA interactors of dArc1 and Copia proteins using RNA immunoprecipitation followed by sequencing (RIP-seq). This screen identified the long non-coding RNA *Stae* as a top candidate that co-immunoprecipitated with both proteins (Xiao et al., 2023; M’Angale et al., 2025). Notably, *Stae* was reported to be upregulated in the aging *Drosophila* brain (Davie et al., 2018), a condition in which retrotransposons are also activated (Li et al., 2013). We then sought to validate these associations using RIP-dPCR. We confirmed a robust and specific association between *Stae* and Copia proteins in S2 cells and body wall muscles (BWMs), but not in the central nervous system (CNS) (Figure 1A). However, we were unable to validate an association between *Stae* and dArc1 (Figure 1A).

**Figure 1.**
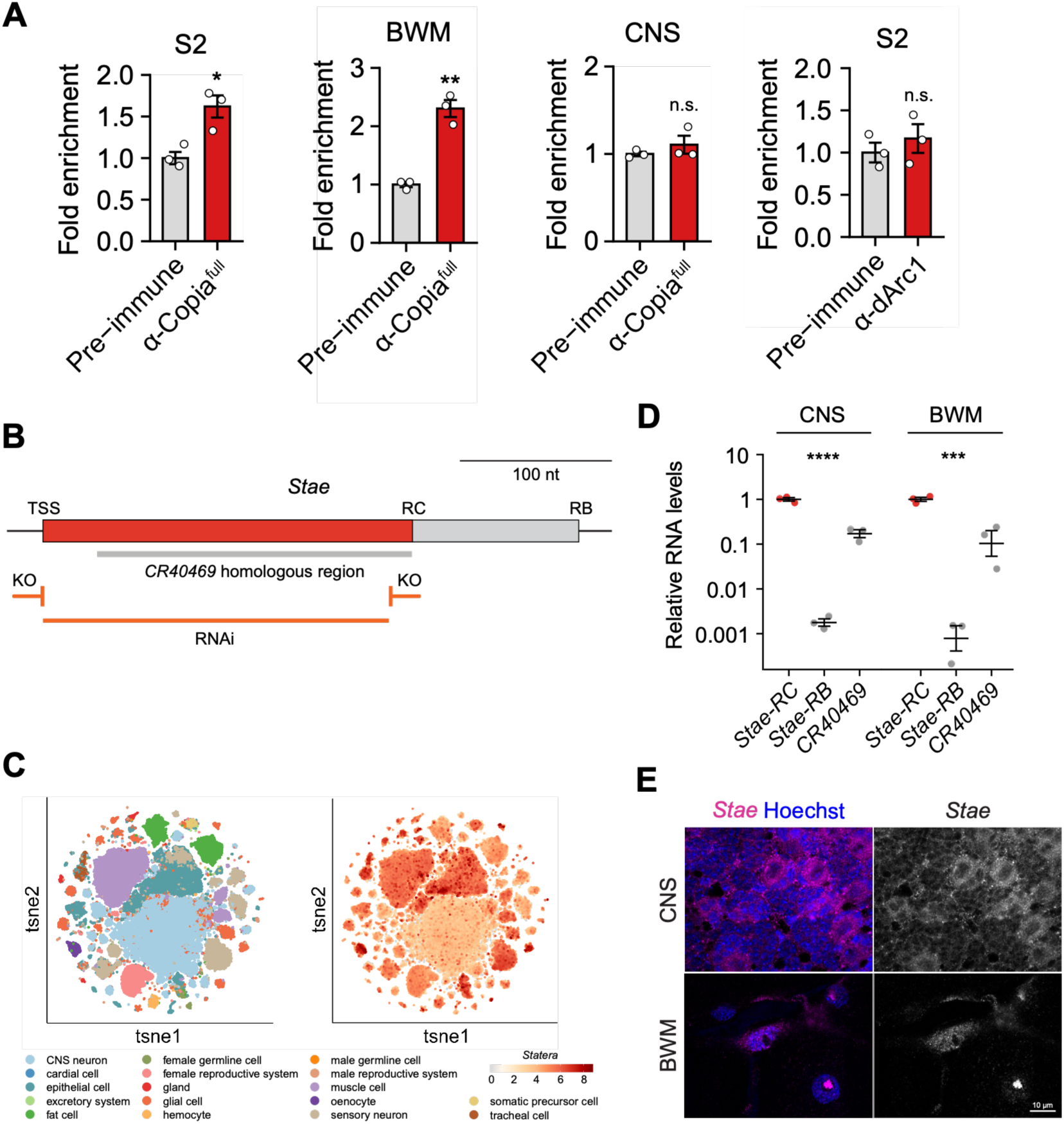
*Stae* is expressed in the CNS and BWMs of *Drosophila* larvae. **(A)** RNA immunoprecipitation using anti-Copia^full^ or anti-dArc1 antibodies. Lysates from *Drosophila* S2 cells, the central nervous system (CNS), or body wall muscles (BWMs) were incubated with antibody-conjugated beads. RNAs were then eluted from beads and detected by *Stae*-specific probes using dPCR. All *Stae* levels were normalized to input. The levels from RIP samples were normalized to control groups. **(B)** Diagram of *Stae* sequences and genetic models used in this study. TSS, transcription start site. RC, the transcription termination site for *Stae*-RC. RB, the transcription termination site for *Stae*-RB. **(C)** *Stae* expression profile in adult *Drosophila*. Data were obtained and visualized using an online single-cell RNA-seq analysis platform (Lu et al., 2023). **(D)** Quantification of expression levels of *Stae* and its homologous sequences in female larval progeny of Canton-S (CS) × *y^1^ w^1118^* flies by dPCR. RNA levels were quantified with target-specific probes and normalized to *rpl32* levels. **(E)** HCR RNA FISH of *Stae* in the larval CNS and BWMs. The localization of *Stae* was detected using *Stae*-specific HCR probes. Nuclei were stained with the Hoechst 33342 DNA dye. Data are presented as mean ± SEM, with each dot representing a biological replicate. Statistical analyses were conducted using Student’s t-test or one-way analysis of variance (ANOVA) followed by Tukey’s post hoc test. ns, not significant; * p<0.05; ** p<0.01; *** p<0.001.

To characterize the *Stae* locus, we examined its genomic organization and expression. The *Stae* gene (*CR34335*) is an intronic lncRNA on the X chromosome with two isoforms and a homolog, *CR40469* (Figure 1B). Analysis of a public single-cell RNA-seq dataset revealed that *Stae* is broadly expressed across *Drosophila* tissues (Lu et al., 2023) (Figure 1C). To further understand *Stae* expression in larval tissues, we quantified the levels of all three *Stae*-related transcripts in the CNS and BWMs by dPCR and found that the *Stae-RC* isoform is by far the most abundant species (Figure 1D). We then used hybridization chain reaction (HCR) RNA FISH to examine the localization of *Stae* RNA and found that it displayed distinct localization patterns in the CNS and BWMs (Figure 1E). In the CNS, *Stae* is diffusely localized in the cytoplasm of nerve cells, whereas in BWMs, its signals are enriched in nucleolus-like structures (Figure 1E). Interestingly, when we attempted to verify the genomic presence of the homolog *CR40469* in our wild-type Canton-S (CS) strain, we discovered that the sequence was absent (Figure S1A and S1B), suggesting it is dispensable under normal laboratory conditions. These findings led us to focus our functional analysis on the *Stae-RC* sequence.

### *Stae* regulates synaptogenesis at the NMJ

To determine the role of *Stae* in synaptogenesis, we analyzed the morphology of the larval neuromuscular junction (NMJ), where synaptic boutons are categorized into distinct subtypes, including type Ib and type Is (Ashley & Budnik, 2017). We generated a *Stae*-null allele using CRISPR and a UAS-*Stae*-RNAi line for tissue-specific KD. Immunostaining of the NMJ in *Stae* KO animals generated by crossing the *Stae*-null allele to a deficiency line revealed a significant decrease in the number of type Is boutons and a reduction in NMJ length, although the total bouton number remained unchanged (Figure 2A and 2B). To determine the tissues in which *Stae* functions, we next performed tissue-specific RNAi knockdown. Expressing *Stae* RNAi with either the pan-neuronal driver *elav-Gal4* or the motor neuron-specific driver *C380-Gal4* caused a significant reduction in total bouton number and led to a more stacked NMJ morphology (Figure 2C, 2D, S1C, and S1D). No significant morphological changes were observed at the NMJ of muscle-specific *Stae* RNAi larvae (data not shown). These results indicate that *Stae* functions presynaptically in motor neurons to promote synaptic growth at the *Drosophila* NMJ.

**Figure 2.**
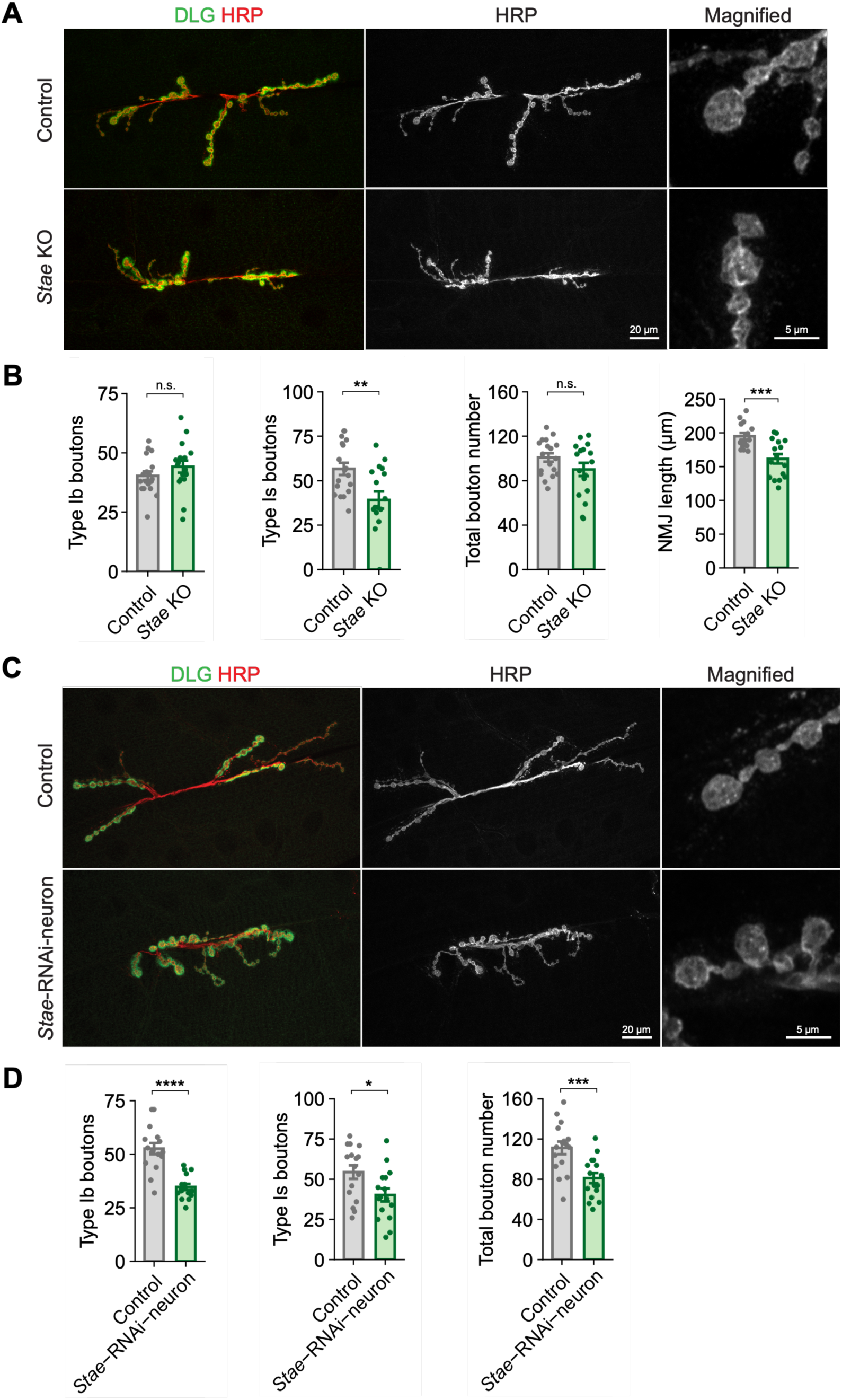
*Stae* KO and neuronal *Stae* RNAi cause a decrease in bouton numbers and altered NMJ morphology. **(A)** Fluorescence images of NMJ preparations from *Stae* KO mutants. Examples of type Ib and type Is boutons are indicated by a white arrow and a white arrowhead, respectively. **(B)** Quantification of NMJ morphology in *Stae* KO. **(C)** Fluorescence images of NMJ preparations from neuronal *Stae* RNAi animals. **(D)** Quantification of NMJ morphology in neuronal *Stae* RNAi. DLG, Discs large (postsynaptic marker); HRP, horseradish peroxidase (presynaptic marker). Data are presented as mean ± SEM, with each dot representing a biological replicate. Statistical analyses were conducted using Student’s t-test. ns, not significant; * p<0.05; ** p<0.01; *** p<0.001.

### *Stae* negatively regulates the retrotransposon *copia*

Given that *Stae* associates with the Copia protein and regulates synaptogenesis, we next sought to determine whether it is involved in the ViSyToR pathway. We first tested the hypothesis that *Stae* is a downstream target of Copia or dArc1. However, when we knocked down either *dArc1* or *copia* in motor neurons, we observed no significant change in *Stae* RNA levels by RT-qPCR, indicating that *Stae* is not regulated by these ViSyToR components at the RNA level (Figure 3A). We then tested the hypothesis that *Stae* is an upstream regulator. To this end, we performed RNA-seq followed by differential expression analysis (DEA) on CNS and BWM tissues from *Stae* KO mutants. Strikingly, *copia* was among the most highly upregulated transposons in the absence of *Stae* (Figure 3B). We validated this result by dPCR and found a significant upregulation of both the *copia^full^*and *copia^gag^* isoforms, while *dArc1* levels remained unchanged (Figure 3C). These data strongly support a model where *Stae* functions as an upstream repressor of *copia*.

**Figure 3.**
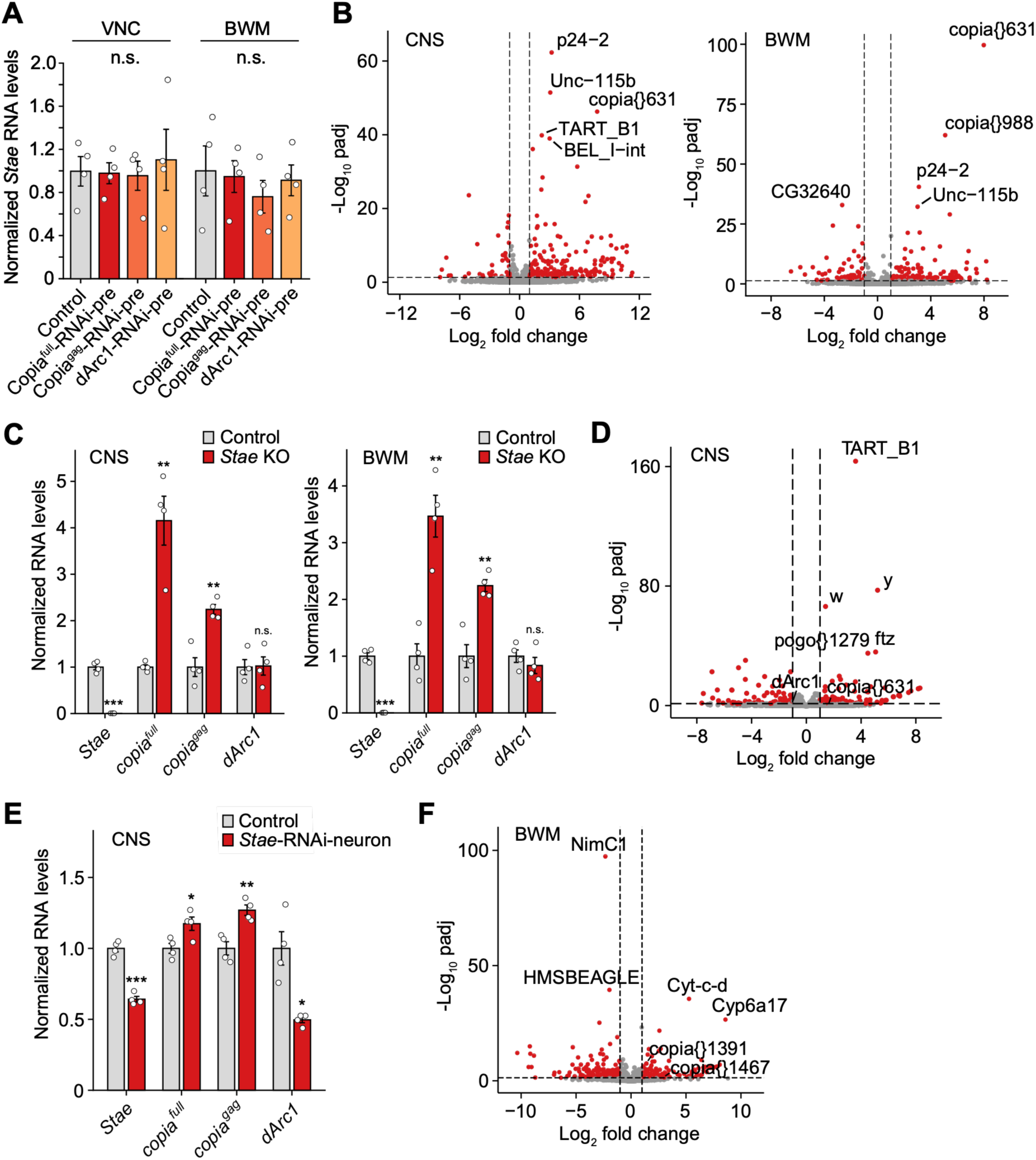
*copia* is upregulated in *Stae* KO and RNAi. **(A)** RT-qPCR quantification of *Stae* levels in Copia and dArc1 RNAi animals. The ventral nerve cord (VNC) and BWMs were dissected from larvae expressing Copia^full^, Copia^gag^, or dArc1 RNAi driven by the motor neuron-specific C380-Gal4 driver. *Stae* RNA levels were quantified and normalized to *rpl32* levels. **(B)** RNA-seq followed by differential expression analysis (DEA) in the CNS and BWMs of *Stae* KO mutants. **(C)** dPCR validation of *copia* and *dArc1* RNA level changes in *Stae* KO. RNA levels were quantified with target-specific probes and normalized to *rpl32* levels. **(D)** RNA-seq followed by differential expression analysis (DEA) in the CNS of neuron-specific *Stae* RNAi larvae. **(E)** dPCR validation of *copia* and *dArc1* RNA level changes in neuron-specific *Stae* RNAi. RNA levels were quantified with target-specific probes and normalized to *rpl32* levels. **(F)** RNA-seq followed by differential expression analysis (DEA) in BWMs of muscle-specific *Stae* RNAi larvae. In the volcano plots, data points meeting the thresholds of adjusted p-value (padj) ≤ 0.05 and |log2 fold change| ≥ 1 are shown in red. In all other graphs, data are presented as mean ± SEM, with each dot representing a biological replicate. Statistical analyses were conducted using one-way ANOVA followed by Tukey’s post hoc test. ns, not significant; * p<0.05; ** p<0.01; *** p<0.001.

We then investigated whether this regulation occurs in specific tissues. In pan-neuronal *Stae* RNAi larvae, differential expression analysis revealed an upregulation of a specific copy, *copia{631}* (Figure 3D), and dPCR confirmed a slight increase in total *copia* RNA levels (Figure 3E). Interestingly, *dArc1* RNA was downregulated in this context, possibly due to competition with derepressed *copia* (Figure 3E). We also performed muscle-specific *Stae* RNAi, which revealed upregulation of a different set of *copia* copies (Figure 3F). Together, these results indicate that while *Stae* acts as a global inhibitor of *copia*, its effects on individual *copia* copies may vary in different tissues.

To have a broader view of *Stae* function, we performed Gene Ontology (GO) analyses on those genes whose expression levels are significantly changed (adjusted p < 0.05) in the CNS of neuronal *Stae* RNAi larvae or BWMs of muscle *Stae* RNAi larvae (Figure S2A). Biological pathways related to protein folding and response to incorrect proteins were significantly enriched in both neuronal and muscle KD, suggesting that *Stae* may also contribute to the maintenance of protein homeostasis. We selected and validated several candidate genes, based on their large fold changes and annotated functions, including *tut*, a gene involved in germ cell development, and the synaptic genes *Unc-115b* and *dpr4* (Figure S2B). The upregulation of *tut* may be related to altered TE expression in *Stae* mutants, given the close link between germline development and TE control.

### *Stae* reduces Copia protein accumulation at the NMJ

Having established that *Stae* represses *copia* RNA, we next asked whether *Stae* also regulates Copia protein levels at the NMJ. We previously showed that Copia protein is enriched at the NMJ and is transferred from presynaptic neurons to postsynaptic muscles (M’Angale et al., 2025). Using immunostaining, we quantified Copia protein levels and found that they were significantly increased at presynaptic sites in both *Stae* KO mutants and neuron-specific *Stae* KD animals (Figure 4A-D). Restricting *Stae* RNAi specifically to motor neurons using the *C380-Gal4* driver was sufficient to recapitulate the increase in presynaptic Copia protein (Figure 4E and 4F). These data demonstrate that *Stae* inhibits Copia protein accumulation at the synapse.

**Figure 4.**
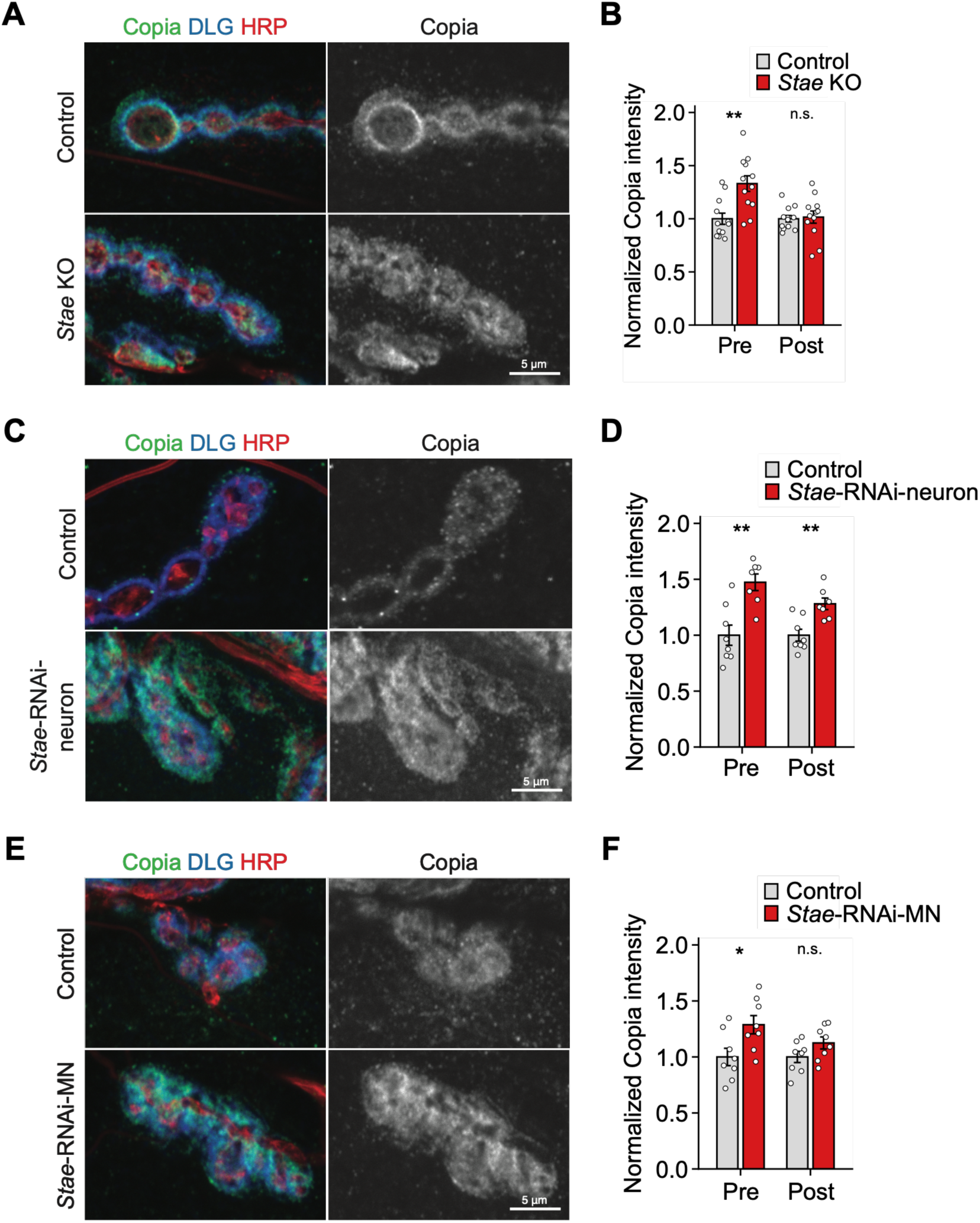
Copia protein levels are increased at the NMJ in *Stae* KO and RNAi larvae. **(A and B)** Copia immunostaining **(A)** and quantification of Copia fluorescence intensity **(B)** at the NMJ in *Stae* KO. **(C and D)** Copia immunostaining **(C)** and quantification of Copia fluorescence intensity **(D)** at the NMJ in neuron-specific *Stae* RNAi. **(E and F)** Copia immunostaining **(E)** and quantification of Copia fluorescence intensity **(F)** at the NMJ in motor neuron-specific *Stae* RNAi. Copia protein was detected with the anti-Copia^full^ antibody. DLG, Discs large (postsynaptic marker); HRP, horseradish peroxidase (presynaptic marker); Pre, presynaptic sites; Post, postsynaptic sites. Copia fluorescence intensities were quantified using Volocity and normalized to controls. Data are presented as mean ± SEM, with each dot representing a biological replicate. Statistical analyses were conducted using Student’s t-test. ns, not significant; * p<0.05; ** p<0.01; *** p<0.001.

### *Stae* promotes synaptogenesis by repressing *copia*

To establish the functional hierarchy between *Stae* and *copia* in the ViSyToR pathway, we performed a genetic rescue experiment to simultaneously knock down *Stae* and *copia* with the *C380-Gal4* driver. Strikingly, *copia* RNAi KD in motor neurons was sufficient to rescue the abnormal NMJ morphology observed in *Stae* RNAi larvae, restoring bouton numbers to control levels (Figure 5A and 5B). These results suggest that the synaptogenesis defects observed in *Stae* RNAi result from *copia* derepression and support a model in which *Stae* promotes synaptogenesis by repressing *copia* in the ViSyToR pathway.

**Figure 5.**
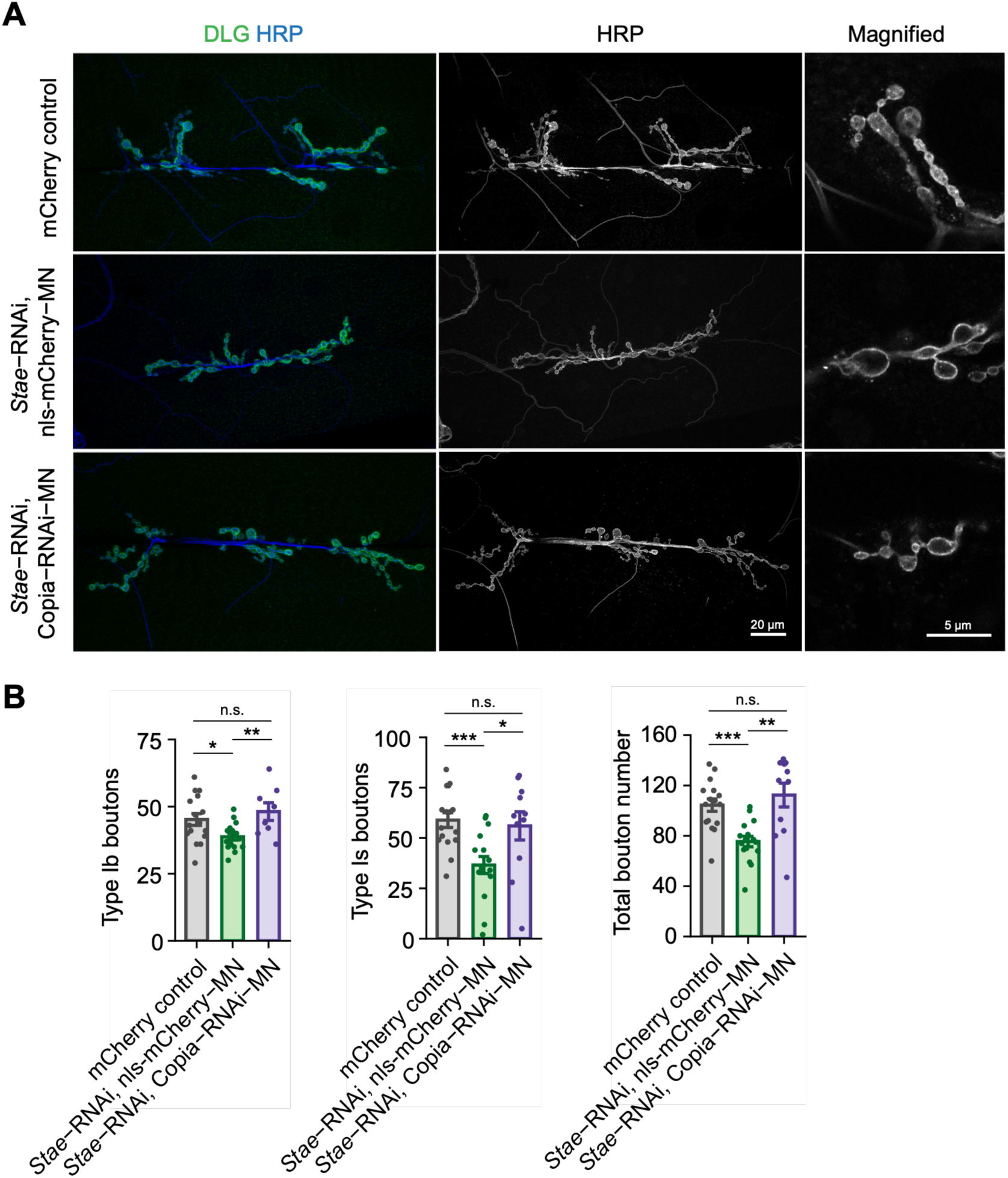
Co-expression of *copia* RNAi in motor neurons rescues the phenotypes observed in *Stae* RNAi. **(A)** Fluorescence images of NMJ preparations from *Stae* and *copia* double RNAi larvae. DLG, Discs large (postsynaptic marker); HRP, horseradish peroxidase (presynaptic marker). **(B)** Quantification of bouton numbers in *Stae* and *copia* double RNAi larvae. Data are presented as mean ± SEM, with each dot representing a biological replicate. Statistical analyses were conducted using one-way ANOVA followed by Tukey’s post hoc test. ns, not significant; * p<0.05; ** p<0.01; *** p<0.001.

### *Stae* reduces *copia* RNA stability and limits *copia* DNA copy number

To determine the mechanism by which *Stae* represses *copia*, we next investigated its role in regulating *copia* RNA stability. We performed an actinomycin D (ActD) chase experiment in CNS and BWM tissues to inhibit transcription and monitor *copia* RNA decay. In control tissues, we observed a transient increase in *copia* levels within the first two hours. Similar responses have been reported for the retrovirus HIV-1 (Cassé et al., 1999; Imamichi et al., 2005). This initial peak was blunted in *Stae* KO tissues, likely due to elevated baseline levels of *copia*. Following this phase, *copia* RNA decayed rapidly in controls, while its decline in *Stae* KO tissues was markedly slower (Figure 6A). In control CNS tissues, we also observed a counterintuitive increase of *copia* levels after 4 hr, possibly reflecting declining tissue viability due to prolonged ActD treatment (Figure 6A). To further quantify *copia* RNA decay, we focused on the decay phase from 2 to 4 hr and found that the stability of *copia* RNA was significantly increased in *Stae* KO tissues (Figure 6B). These data suggest that *Stae* reduces *copia* RNA stability.

**Figure 6.**
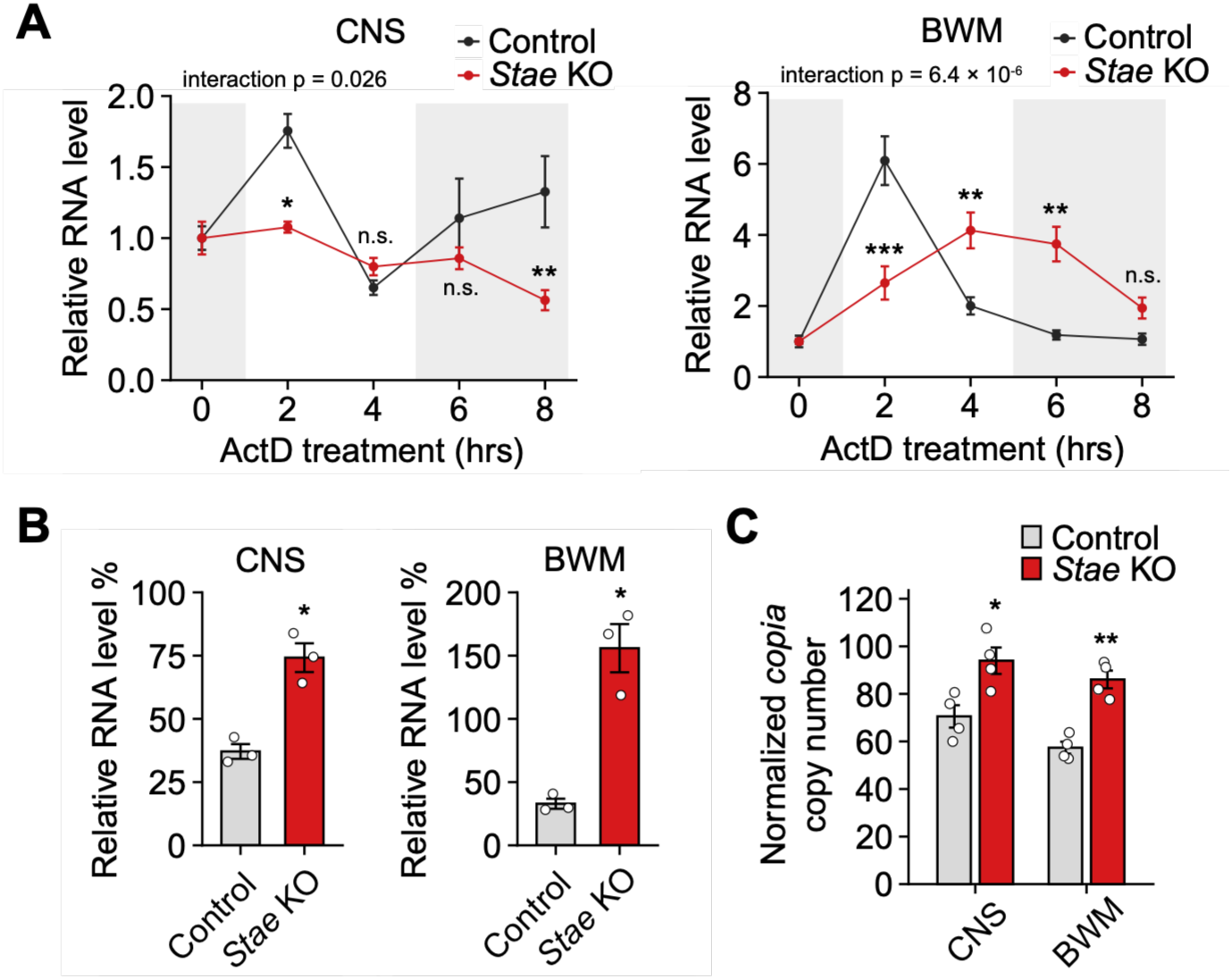
*Stae* KO mutants show an increase in *copia* RNA stability and DNA copy number. **(A)** RNA stability assay of *copia* in *Stae* KO mutants. The CNS and BWMs of *Drosophila* larvae were dissected and incubated with S2 media supplemented with actinomycin D (135 μM), a transcription inhibitor. Samples were then collected at individual time points, and RNA was extracted. *copia* levels were quantified using *copia^full^*-specific probes and normalized to 18S rRNA levels, which remain relatively stable during the treatment. **(B)** Ratio of *copia* RNA level at 4 hr to 2 hr during ActD treatment. The *copia* RNA levels at 4 hr in (A) were normalized to the RNA levels at 2 hr and expressed as percentages of the 2 hr values. **(C)** Copy number analysis of *copia* in *Stae* KO mutants. The CNS and BWMs of *Drosophila* larvae were dissected, and DNA was extracted from these tissues. *copia* DNA levels were determined using *copia* LTR-specific probes and normalized to the DNA levels of the *actin* promoter. Data are presented as mean ± SEM, with each dot representing a biological replicate. n = 3 biological replicates in (A). In (A), statistical analyses were conducted using two-way ANOVA followed by Bonferroni-corrected pairwise comparisons at each time point. CNS (left), time *p* = 0.0021, genotype *p* = 0.0016, interaction (time × genotype) *p* = 0.026. BWM (right), time *p* = 3.3 × 10^-5^, genotype *p* = 0.089, interaction *p* = 6.4 × 10^-6^. In (B) and (C), statistical analyses were conducted using Student’s t-test. ns, not significant; * p<0.05; ** p<0.01; *** p<0.001.

Since *Stae* represses *copia* expression, we asked whether it affects *copia* DNA copy number, which can increase as a result of retrotransposition. While *copia* is generally inactive in somatic cells, its retrotransposition can occur under certain conditions (Pasyukova & Nuzhdin, 1993; Pasyukova et al., 1997). We used dPCR to quantify the DNA copy number of *copia* LTRs in both the CNS and BWMs and observed a significant increase in *copia* copy numbers in *Stae* KO mutants (Figure 6C). This increase may result from two non-exclusive scenarios: first, loss of *Stae* leads to somatic transposition in the CNS and BWMs during larval development, a rare but reported phenomenon in *Drosophila* neuronal lineages (Di Franco et al., 1989; Treiber & Waddel, 2017; Siudeja et al., 2021). Second, transpositions could occur and accumulate over generations in the homozygous *Stae*-null stock line, a process documented for *copia* (Pasyukova & Nuzhdin, 1993; Pasyukova et al., 1997). Future experiments will be needed to distinguish between these possibilities. Overall, these data provide evidence that *Stae* limits the accumulation of *copia* DNA copies, potentially by repressing *copia* retrotransposition.

## DISCUSSION

Our investigation into the regulation of the ViSyToR pathway began with the identification of the lncRNA *Stae* as an interactor of Copia protein, which was later validated by RIP-dPCR in S2 cells and BWMs. The dPCR analyses revealed that *Stae* is expressed at a remarkably high level, approximately 5-10 times that of the housekeeping gene *rpl32*. The high abundance of *Stae*, in contrast to the typically low expression of lncRNAs, prompted us to study its potential physiological role (Derrien et al., 2012).

Our functional analysis confirms that *Stae* is a positive regulator of synaptogenesis in *Drosophila*. The phenotypes observed upon disrupting *Stae* expression, including a general decrease in bouton number, reduced NMJ length, and a more stacked bouton layout (Figure 2), are the inverse of previously reported *copia* KD phenotypes (M’Angale et al., 2025). The decrease in type Is boutons in *Stae* KO animals is also the opposite of what we observed with *copia* KD (Figure 2B) (M’Angale et al., 2025). *Stae* KO mutants also displayed a high degree of phenotypic variability, which has been reported during TE derepression as a result of genomic instability (Niu et al., 2019; Latzel et al., 2023). Consistent with opposite phenotypes of *Stae* and *copia* disruption, we showed that *Stae* is an upstream repressor of *copia*. The regulation of *copia* by *Stae* was observed in both the CNS and BWMs. However, different *copia* copies were preferentially affected (Figure 3D and 3F), possibly reflecting tissue-specific variations in local chromatin states or transcript abundance (Pasyukova et al., 1997). In addition, *Stae* inhibits the enrichment of Copia protein at presynaptic sites, supporting a model in which *Stae* promotes synaptic growth by reducing Copia protein accumulation.

One interesting observation in our study is the discrepancy between the phenotypes of the *Stae* KO and neuronal KD models. The increase in *copia* RNA levels correlated well with the extent of *Stae* depletion (Figure 3C and 3E), while the degree of Copia protein accumulation and the severity of the NMJ phenotype did not (Figures 2 and 4). The neuronal KD produced a generally stronger synaptic phenotype than the whole-animal KO, despite a more modest increase in *copia* RNA. The mild phenotype in the KO model, despite high *copia* RNA levels, may point to the activation of developmental compensatory mechanisms. It is plausible, for example, that a whole-animal knockout triggers systemic feedback loops that limit Copia protein production, thereby buffering its effect at the synapse. Conversely, the stronger phenotype in the RNAi model could be attributed to the RNAi construct targeting both *Stae* and its homolog *CR40469*. However, this is less likely given that our wild-type flies lacking *CR40469* have normal NMJs. Interestingly, a recent study showed that *CR40469* is upregulated during *Drosophila* wing regeneration and is responsible for wing regeneration (Camilleri-Robles et al., 2024). The ectopic expression of *Stae* partially rescued the wing phenotype observed in *CR40469* mutants (Camilleri-Robles et al., 2024). We speculate that *Stae* and *CR40469* might share overlapping functions related to cell growth regulation, but act under different conditions in *Drosophila*.

Another important question is where *Stae* executes its biological functions. Given that *copia* is upregulated in both CNS and BWM tissues of *Stae* KO larvae, the *copia* repression mediated by *Stae* is likely a universal mechanism in *Drosophila*. It would be interesting to test if *Stae* has similar functions in germline tissues such as the ovary and testis, which are active sites for transposon repression. However, a pronounced NMJ phenotype was observed upon neuronal *Stae* RNAi, whereas we did not observe any significant changes in NMJ morphology in muscle-specific *Stae* RNAi larvae (data not shown). Thus, we propose that *Stae* mainly functions in neurons to regulate synapse formation. Consistent with this, we previously reported that *copia* KD in motor neurons leads to bouton overgrowth (M’Angale et al., 2025). We also discussed the possibility of a non-cell-autonomous mechanism, where *copia* elements are transferred from presynaptic neurons to postsynaptic muscles and inserted into the genome (M’Angale et al., 2025), though the data collected so far were insufficient to support this hypothesis.

The detailed mechanisms by which *Stae* represses *copia* remain to be elucidated. Our data support a post-transcriptional model in which *Stae* destabilizes *copia* RNA. In *Drosophila*, the piRNA pathway mediated by PIWI proteins represents a primary mechanism for TE repression, especially in the germline (Brennecke et al., 2007; Ai et al., 2009). The endo-siRNA pathway has also been reported to contribute to TE repression, mainly in somatic tissues (Gu et al., 2009; Stein et al., 2015). Both *copia* piRNAs and siRNAs have been detected in *Drosophila* cultured cells, suggesting that *copia* expression might be regulated by both pathways (Chung et al., 2008). Upregulation of *copia* RNA levels has been observed in the ovaries of *Drosophila* Piwi mutants (Wang et al., 2015) and *copia* piRNAs were found to be associated with all three PIWI proteins (Klumpe et al., 2025). The potential roles of lncRNAs in TE repression have been reported but mechanistic studies are still in their infancy. Several studies have shown that lncRNAs can repress TE expression by interacting with TE RNA or DNA and inducing chromatin modifications (Berretta et al., 2008; Zhang et al., 2022). To our knowledge, regulation of TE RNA stability by a lncRNA has not been reported. Although we cannot rule out the possibility that *Stae* regulates *copia* chromatin states, our data suggest an alternative mechanism in which a lncRNA represses the expression of a TE by affecting its RNA stability. In the future it would be interesting to examine the effects of *Stae* on the piRNA or siRNA pathways in *Drosophila*. In our RIP-dPCR experiments, we showed that *Stae* associates with Copia proteins in S2 cells and BWMs but not in the CNS, suggesting that the association between *Stae* and Copia proteins might not be required for *Stae* function in neurons, but might instead be a byproduct of *Stae*-mediated *copia* regulation. One compelling hypothesis is that *Stae* acts as a molecular scaffold, binding to both the *copia* transcript and components of the RNA decay machinery to facilitate its degradation. Combining previous results and current lines of evidence, we propose a model in which *Stae* represses the expression of the retrotransposon *copia*, a negative regulator of synaptogenesis, by destabilizing its transcripts. As a result, *Stae* reduces *copia* RNA levels, DNA copy number, and Copia protein accumulation in neurons, and thereby promotes proper synaptic growth at the NMJ (Figure 7).

**Figure 7.**
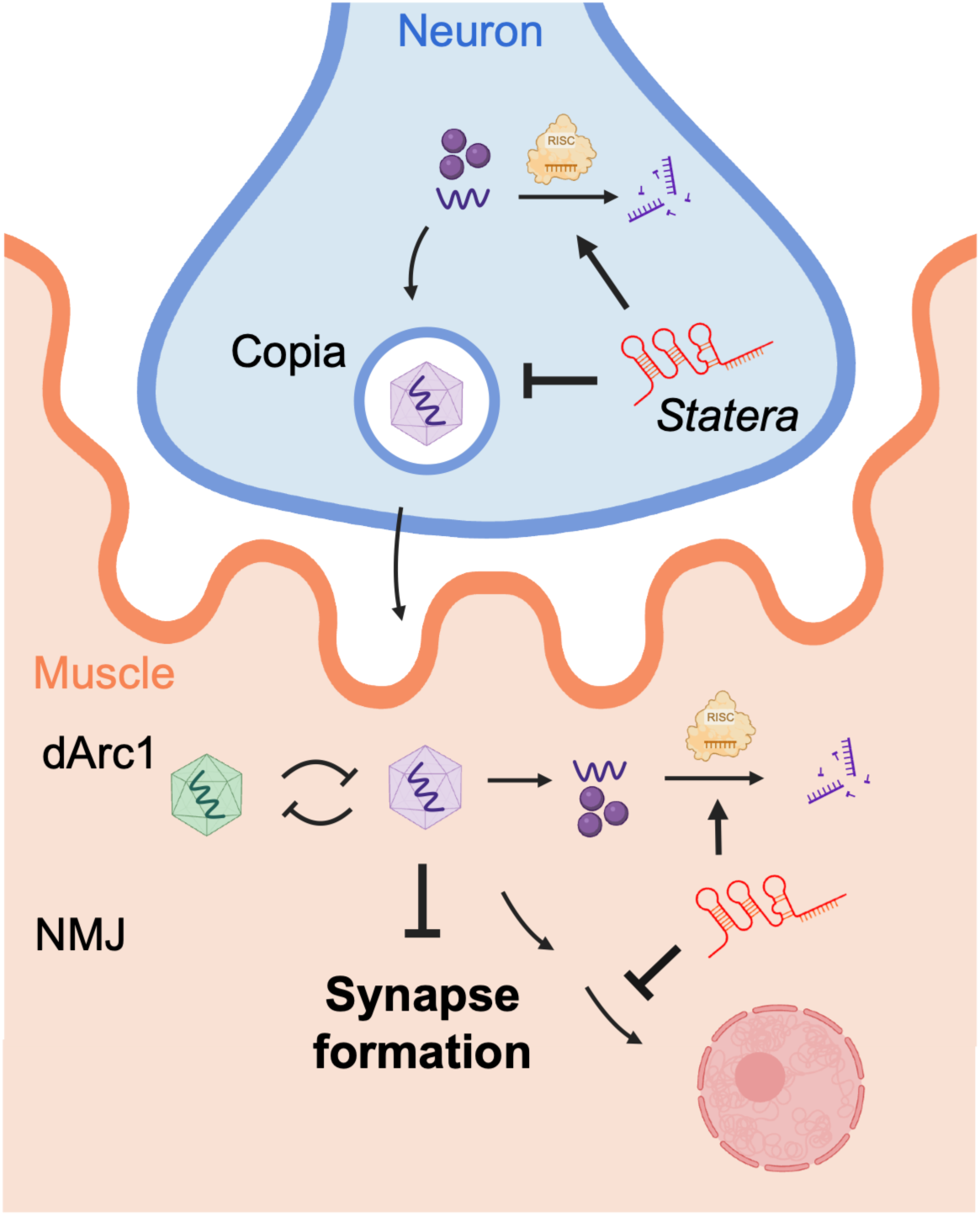
Current model of *Stae* function at the *Drosophila* larval NMJ. *Stae* functions as a negative regulator of *copia* in the ViSyToR pathway. *Stae* reduces *copia* RNA stability, limits *copia* DNA copy number, and decreases Copia protein levels, thereby promoting synapse formation at the NMJ.

## METHODS

### Experimental models

The following fly lines were used: Canton-S (BDSC 1), y[1] w[1118];; attP2{nos-Cas9}/TM6C,Sb Tb (Bestgene), y[1] w[1118], *Stae*-null (see below), y[1] w[1118] (see below), Df(1)ED6712/FM7a-chFP (see below), UAS-Copia^full^-RNAi (M’Angale et al., 2025), UAS-Copia^gag^-RNAi (M’Angale et al., 2025), UAS-dArc1-RNAi (Ashley et al., 2018), UAS-*Stae*-RNAi (see below), UAS-mCherry-NLS (BDSC 38424), elav-Gal4 (BDSC 8765), C380-Gal4 (Budnik, 1996), C57-Gal4 (Budnik, 1996).

### Fly husbandry and S2 cell culture

All flies were raised on standard molasses formulation food at either 25 °C (KO crosses and other crosses) or 29 °C (RNAi crosses).

S2 cells were cultured at 25 °C in Schneider’s *Drosophila* Medium (Gibco) supplemented with 10% fetal bovine serum (Hyclone), 50 unit/ml penicillin, and 0.05 mg/ml streptomycin solution (Sigma).

### Generation of a *Stae*-null allele

For pCFD5_w-StaeKO, the two single-guide RNA (sgRNA) sites were selected by maximizing the on-target score and minimizing the off-target effects using Benchling (https://www.benchling.com/) and CRISPR target finder (http://targetfinder.flycrispr.neuro.brown.edu/). The gRNA inserts were synthesized and cloned into the pCFD5_w plasmid (Addgene 112645) (Port et al., 2014) by GenScript.

A mutant allele of *Statera* was generated by CRISPR/Cas9-mediated genomic engineering. The pCFD5_w-StaeKO construct was injected into Cas9 expressing embryos (y[1] w[1118];; attP2{nos-Cas9}/TM6C,Sb Tb from BestGene). The resulting adults were crossed to a FM7c balancer fly line (BDSC 36337). The *Stae*-null mutants were selected by PCR using genotyping primers (Stae-gF,R) and made into stocks. For *Stae* KO, the *Stae*-null males were crossed to a deficiency line Df(1)ED6712/FM7a-chFP (BDSC 9169, rebalanced on FM7a-chFP) to minimize effects of potential off-target mutations. The third-instar larvae with no red fluorescence were selected (*Stae*-null/Df(1)ED6712). For the control, the y[1] w[1118];; attP2{nos-Cas9}/TM6C,Sb Tb flies were balanced on the same FM7c balancer chromosome and nos-Cas9(III-attP2) were removed by selecting Sb adults. The TM6C,Sb Tb chromosome was then removed by self-crossing and made into stocks. The resulting y[1] w[1118] males were crossed to Canton-S and the third-instar larvae were used for controls.

### Generation of a *Stae* RNAi line

The *Stae*-RNAi insert (long dsRNA) for *Stae* was synthesized by GenScript and cloned into pWALIUM10-roe (Perkins et al., 2015) following a TRiP protocol (https://fgr.hms.harvard.edu/sites/g/files/omnuum5366/files/fly/files/ko_protocol.pdf). The construct was injected into flies and insertion was accomplished using phiC-31 integrase by Bestgene. The resulting UAS responder line was then crossed with various Gal4 driver lines to achieve tissue-specific *Stae* knockdown. The constructs were then injected into flies and insertion was accomplished using phiC-31 integrase by Bestgene.

### Immunocytochemistry

Third instar larval body wall muscles (BWMs) were dissected in calcium-free saline and fixed in either Bouin’s fixative (for anti-Copia^full^ staining) or 4% (w/v) paraformaldehyde in 0.1 M PBS (for all other staining). Fixed samples were then washed and permeabilized in PBST (0.1 M PBS, 0.2% (v/v) Triton X-100) 3 times for 10 min each, and incubated with primary antibodies overnight at 4°C. On the second day, the samples were washed 3 times with PBST, incubated with secondary antibodies for 1-2 h at room temperature, washed again 3 times with PBST, and mounted in Vectashield Hardset Mounting Media (Vector Laboratories Inc.). The primary antibodies were used: rabbit anti-DLG, 1:40,000 (Koh et al., 1999), mouse anti-DLG, 1:200 (DSHB 4F3), rabbit anti-Copia^full^, 1:1,000 (M’Angale et al., 2025). The secondary antibodies were obtained from Jackson ImmunoResearch: Alexa Fluor-594-conjugated goat anti-HRP, 1:500, DyLight-405-conjugated goat anti-HRP, Alexa Fluor-488-conjugated donkey anti-Rabbit, Alexa Fluor-594-conjugated goat anti-Rabbit, Alexa Fluor-647-conjugated goat anti-Mouse, all at 1:200 dilution.

### Fluorescent in situ hybridization (FISH)

*Drosophila* third instar BWMs attached with the intact CNS were dissected in calcium-free saline and fixed in fixation solution (PBS, 0.3% Triton X-100, 4% paraformaldehyde) by gentle rocking at room temperature for 45 min. Samples were then washed 3 times with PBTX, transferred to 0.2 mL 70% ice-cold ethanol, and incubated overnight at 4°C.

Given the limited length of *Stae* sequence, hybridization chain reaction (HCR) technology from Molecular Instruments was used for RNA FISH. An HCR probe set consisting of 5 probes was designed for the lncRNA *Stae*. Samples were then hybridized with the HCR probes and signals were amplified by hairpins h1 and h2, following HCR ‘Sample in Solution’ protocol.

On the last day, samples were incubated with Hoechst 33342 for 15 min, washed 3 times, 5 min each in the wash solution (2x SSC with 10% deionized formamide), rinsed with PBS, and mounted in Vectashield Hardset Mounting Media.

### Confocal microscopy and quantification

Z-stacked images (for NMJ arbors) and single-plane images (for magnified boutons) were acquired using a Zeiss LSM 800 confocal microscope equipped with a Zeiss 63X Plan-Apochromat 1.40 NA DIC M27 oil immersion objective and a Zeiss 40X Plan-Apochromat 1.30 NA DIC (UV) VIS-IR M27 oil immersion objective. For NMJ arbors, 2D images were obtained from maximum-intensity projection of Z-stack images. For magnified boutons, single-plane images were captured with higher magnification and lower scanning speed. Images were acquired with identical acquisition settings for samples and corresponding controls. Bouton numbers were counted manually and NMJ lengths (longest branch lengths) were quantified by a previously described Fiji plugin in ImageJ (Castells-Nobau et al., 2017). Fluorescence signals were quantified by Volocity as follows. For presynaptic signals, the bouton volume bounded by HRP staining was selected and fluorescence intensity inside was measured. For postsynaptic signals, the HRP containing volume was subtracted from the DLG containing volume and the intensity within the remaining volume was measured. Fluorescence intensity was determined as the sum of total pixel intensity normalized to bouton volume, and data were normalized to control values.

### Digital PCR (dPCR)

Samples were homogenized in RLT buffer (QIAGEN) with 1:100 diluted β-mercaptoethanol (β-ME). RNA was extracted with RNeasy Micro or Mini Kit (QIAGEN). RNA samples were reverse transcribed into cDNA using Superscript IV first-strand synthesis reaction (Invitrogen) or AzuraQuant II cDNA Synthesis Kit (Azura Genomics) following manufacturer protocols. The dPCRs were multiplexed in 26K 24-well or 8.5K 96-well QIAcuity nanoplates using the QIAcuity system (QIAGEN). Either QIAcuity EvaGreen master mix (QIAGEN) with the gene-specific primer sets (IDT) or probe master mix (QIAGEN) with the gene-specific probe sets (IDT) was used for the reactions. Absolute concentrations (copies/μL) were obtained from the QIAcuity Software Suite (QIAGEN) and normalized to the internal controls.

### RNA stability assay

Larval tissues were dissected and incubated in 250 μL pre-warmed Schneider’s Drosophila Medium (Gibco) with 135 μM actinomycin D (Sigma-Aldrich) in RNase-free 1.5 mL microfuge tubes. Samples were incubated at 25 °C with shaking at 300 rpm and mixed by pipetting every hour. At each time point samples were collected, washed in cold RNase-free PBS, and lysed in RLT buffer (QIAGEN). RNA was extracted with RNeasy Micro or Mini Kit (QIAGEN). RNA samples were reverse transcribed into cDNA using AzuraQuant II cDNA Synthesis Kit (Azura Genomics) following the manufacturer protocol with RNase H digest. Target cDNA concentrations were determined by dPCR using the QIAcuity Software Suite (QIAGEN) and normalized to cDNA level of 18S rRNA.

### Copy number analysis

The *Drosophila* CNS and BWM tissues were dissected and lysed in 180 μL Buffer ATL (QIAGEN) with 20 μL Proteinase K (QIAGEN) at 56°C for 1h. DNA was extracted with QIAamp DNA Mini Kit (QIAGEN). For fragmentation and RNA removal, purified DNA samples were incubated with AluI enzyme (NEB) and 100 µg/mL RNase A (QIAGEN) in 1x rCutSmart buffer (NEB) at 37°C for 1h. DNA fragments were further purified using QIAquick PCR Purification Kit (QIAGEN). The dPCRs were multiplexed in 26K 24-well or 8.5K 96-well QIAcuity nanoplates using the QIAcuity system (QIAGEN). Either QIAcuity EvaGreen master mix (QIAGEN) with the gene-specific primer sets (IDT) or probe master mix (QIAGEN) with the gene-specific probe sets (IDT) was used for the reactions. Target concentrations were obtained from the QIAcuity Software Suite (QIAGEN) and normalized to DNA levels of *actin* promoter.

### RNA sequencing with DEA

RNA was isolated as described previously. RNA concentration and integrity were assessed by Qubit 4 Fluorometer (Thermo Fisher Scientific) and 2100 Bioanalyzer (Agilent), respectively. Samples were then sent to Novogene for RNA-seq, with library preparation using poly(A) enrichment and sequencing using the NovaSeq PE150 platform. Reads were mapped to the *Drosophila* genome by STAR. The transcript expression combining both genes and transposons was quantified using TEtranscripts, and the differential expression genes were explored by DESeq2. GO analysis was performed using clusterProfiler.

### RNA immunoprecipitation

For S2 cell preparations, cells were centrifuged at 1000 g to pellet the cells, washed with ice-cold PBS, and centrifuged again. Pellets were then resuspended in 1 mL ice-cold RIPA (Abcam) lysis buffer supplemented with 1x protease inhibitors (Roche) and 1 U/μL RNase inhibitor (Invitrogen), mixed for 30 min at 4°C, and centrifuged for 20 min at 12,000 rpm at 4°C. Supernatants were transferred to a fresh tube on ice.

For tissue preparation, tissues (10x BWMs and 20x CNS for each biological replicate) were dissected and homogenized in 500 μL ice cold RIPA lysis buffer supplemented with 1x protease inhibitors (Roche) and 1 U/μL RNase inhibitor (Invitrogen) using an electric homogenizer, mixed for 2 h at 4°C, and centrifuged for 20 min at 12,000 rpm at 4°C.

Bead preparation and Immunoprecipitation were performed according to the manufacturer protocol of Protein A/G Magnetic Beads (Pierce). In brief, the resulting supernatants were first pre-cleared with pre-immune serum and beads, and then incubated overnight at 4°C with either 10 μg rabbit anti-Copia^full^ (M’Angale et al., 2025), rabbit anti-dArc1 (Ashley et al., 2018), or an equal amount of pre-immune serum. On the second day, samples were then incubated for 1 h at 4°C with protein A/G magnetic beads and washed 5 times with RIPA buffer. For immunoblotting, beads were incubated directly with 4× protein loading buffer (Li-Cor) with β-ME (Sigma). For digital PCR, RNA was eluted from the beads with RLT buffer (QIAGEN) supplemented with β-ME and then purified using the RNeasy mini kit (QIAGEN) with DNase digest using RNase-free DNase set (QIAGEN).

### Quantification and statistical analysis

Experiments were performed with at least three biological replicates. For comparisons of a single experimental group with control, Student’s t-test was used. For comparisons among multiple experimental groups, a one-way analysis of variance (ANOVA) followed by Tukey’s post hoc test was used. For time-course experiments, two-way ANOVA followed by Bonferroni-corrected pairwise comparisons was used. Statistical analyses were performed using R. Data in graphs are presented as mean ± SEM. Significance levels are presented as, ns, not significant, ∗ p<0.05, ∗∗ p<0.01, and ∗∗∗ p<0.001.

## ACKNOWLEDGEMENTS

We thank members of the Thomson laboratory for helpful discussions and technical support. We also thank the Bloomington Drosophila Stock Center for providing fly stocks and FlyBase for genetic resources. This work was supported by NIH Grant R01NS112492 to T.T.

---

**Figure S1.**
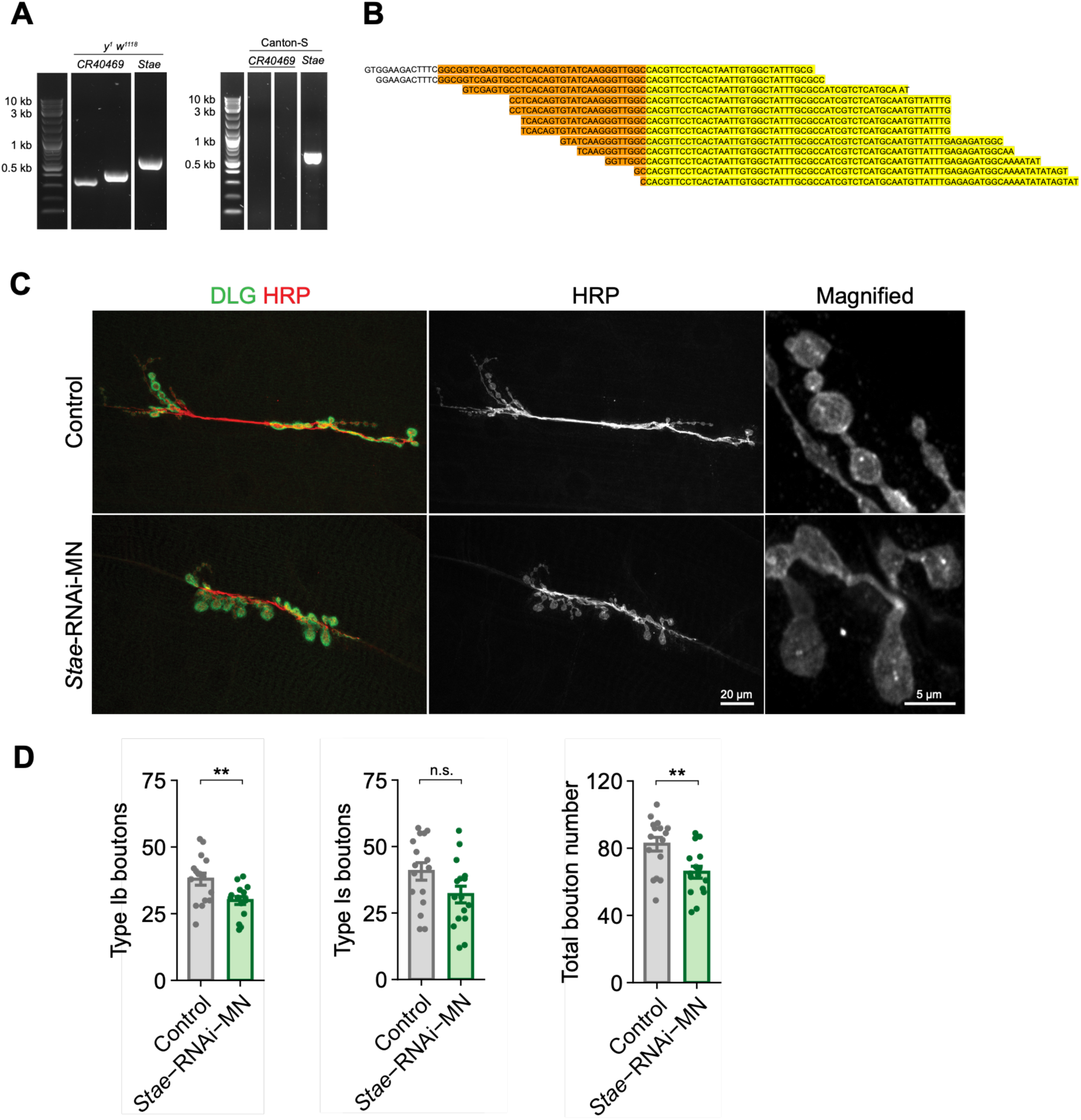
Motor neuron-specific *Stae* RNAi causes altered NMJ morphology. **(A)** DNA gel images show that *Stae* is present in both wild-type CS and *y^1^ w^1118^* flies from BestGene, while the *CR40469* homologous sequence is not detected in the lab CS strain. **(B)** Reads from whole-genome sequencing show that *Stae* fragments, but not *CR40469* fragments, are detected in the lab CS strain. Sequences matching the annotated 5’ region of *CR40469* are highlighted in yellow. Sequences upstream of the annotated *CR40469* sequence are highlighted in orange. These upstream sequences match the 5’ region of *Statera*, but not the genomic sequence upstream of *CR40469*. **(C)** Fluorescence images of NMJ preparations from motor neuron-specific *Stae* RNAi animals. **(D)** Quantification of NMJ morphology in motor neuron-specific *Stae* RNAi. DLG, Discs large (postsynaptic marker); HRP, horseradish peroxidase (presynaptic marker). Data are presented as mean ± SEM, with each dot representing a biological replicate. Statistical analyses were conducted using Student’s t-test. ns, not significant; * p<0.05; ** p<0.01; *** p<0.001.

**Figure S2.**
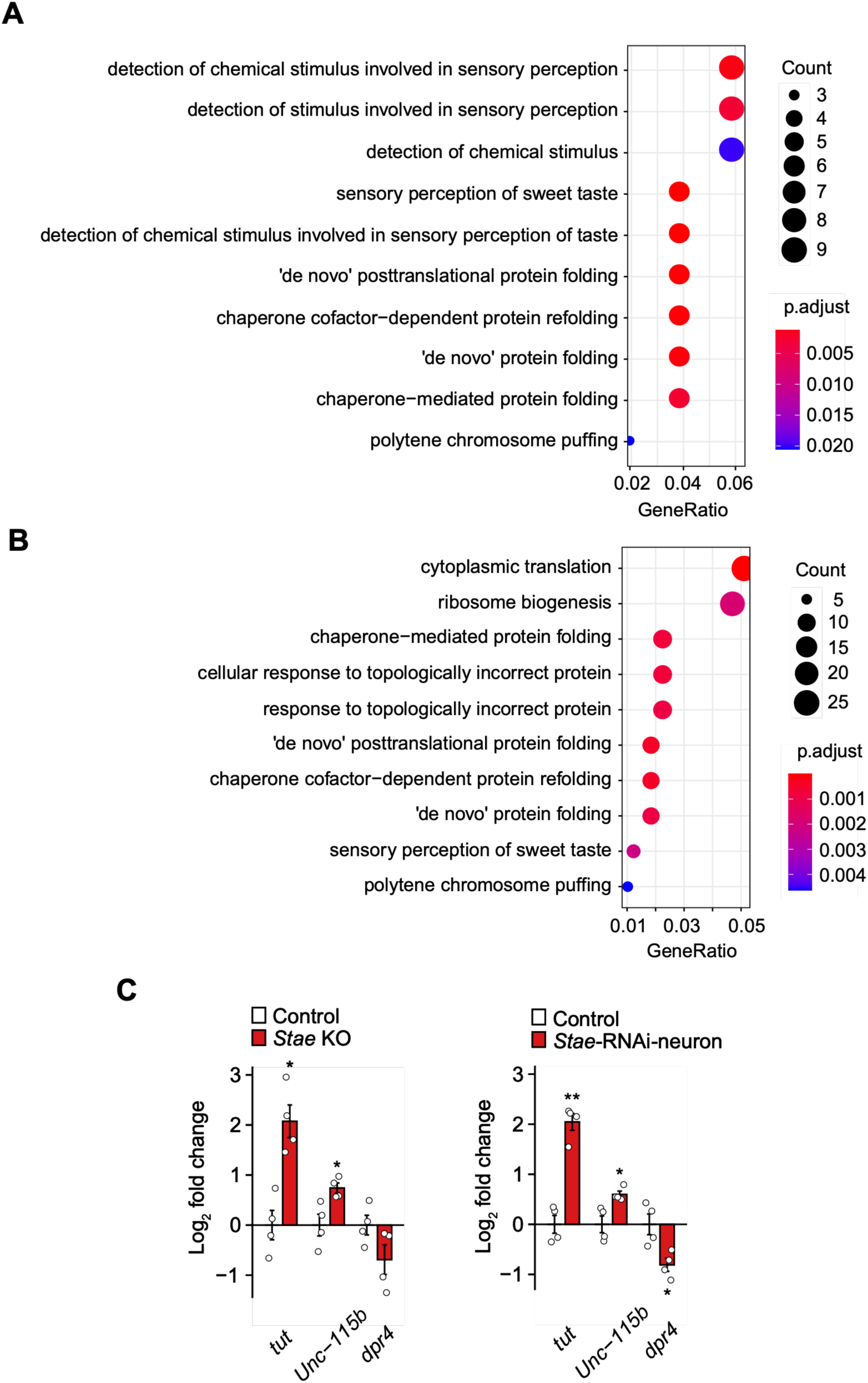
Biological processes associated with *Stae*. **(A and B)** GO analysis of differentially expressed genes in **(A)** neuron-specific *Stae* RNAi and **(B)** muscle-specific *Stae* RNAi. The analyses identified common biological processes related to protein folding, as well as tissue-specific processes such as sensory perception in the CNS and cytoplasmic translation in BWMs. **(C)** dPCR validation of candidate genes downstream of *Stae* in *Stae* KO and RNAi larvae. Total RNA was extracted from the CNS, and the candidate RNA levels were determined using target-specific primer sets and normalized to *rpl32* levels. Data are presented as mean ± SEM, with each dot representing a biological replicate. Statistical analyses were conducted using Student’s t-test. ns, not significant; * p<0.05; ** p<0.01; *** p<0.001.

## REFERENCES

Ai, K. L., Tao, L., & Kai, T. (2009). piRNAs mediate posttranscriptional retroelement silencing and localization to pi-bodies in the Drosophila germline. Journal of Cell Biology, 186(3), 333–342. 10.1083/jcb.200904063

Ashley, J., & Budnik, V. (2017). A Tale of Two Inputs. Neuron, 93(6), 1245–1247. 10.1016/j.neuron.2017.03.013

Ashley, J., Cordy, B., Lucia, D., Fradkin, L. G., Budnik, V., & Thomson, T. (2018). Retrovirus-like Gag Protein Arc1 Binds RNA and Traffics across Synaptic Boutons. Cell, 172(1–2), 262–270.e11. 10.1016/j.cell.2017.12.022

Baker, B. M., Mokashi, S. S., Shankar, V., Hatfield, J. S., Hannah, R. C., MacKay, T. F. C., & Anholt, R. R. H. (2021). The Drosophila brain on cocaine at single-cell resolution. Genome Research, 31(10), 1927–1937. 10.1101/gr.268037.120

Barry, G., Briggs, J. A., Vanichkina, D. P., Poth, E. M., Beveridge, N. J., Ratnu, V. S., Nayler, S. P., Nones, K., Hu, J., Bredy, T. W., Nakagawa, S., Rigo, F., Taft, R. J., Cairns, M. J., Blackshaw, S., Wolvetang, E. J., & Mattick, J. S. (2014). The long non-coding RNA Gomafu is acutely regulated in response to neuronal activation and involved in schizophrenia-associated alternative splicing. Molecular Psychiatry, 19(4), 486–494. 10.1038/mp.2013.45

Bernard, D., Prasanth, K. V., Tripathi, V., Colasse, S., Nakamura, T., Xuan, Z., Zhang, M. Q., Sedel, F., Jourdren, L., Coulpier, F., Triller, A., Spector, D. L., & Bessis, A. (2010). A long nuclear-retained non-coding RNA regulates synaptogenesis by modulating gene expression. The EMBO Journal, 29(18), 3082–3093. 10.1038/emboj.2010.199

Berretta, J., Pinskaya, M., & Morillon, A. (2008). A cryptic unstable transcript mediates transcriptional trans-silencing of the Ty1 retrotransposon in S. cerevisiae. Genes & Development, 22(5), 615–626. 10.1101/gad.458008

Bodea, G. O., Botto, J. M., Ferreiro, M. E., Sanchez-Luque, F. J., de Los Rios Barreda, J., Rasmussen, J., … & Faulkner, G. J. (2024). LINE-1 retrotransposons contribute to mouse PV interneuron development. Nature Neuroscience, 27(7), 1274–1284. 10.1038/s41593-024-01650-2

Bourque, G., Burns, K. H., Gehring, M., Gorbunova, V., Seluanov, A., Hammell, M., Imbeault, M., Izsvák, Z., Levin, H. L., Macfarlan, T. S., Mager, D. L., & Feschotte, C. (2018). Ten things you should know about transposable elements. Genome Biology, 19(1), 199. 10.1186/s13059-018-1577-z

Brennecke, J., Aravin, A. A., Stark, A., Dus, M., Kellis, M., Sachidanandam, R., & Hannon, G. J. (2007). Discrete Small RNA-Generating Loci as Master Regulators of Transposon Activity in Drosophila. Cell, 128(6), 1089–1103. 10.1016/j.cell.2007.01.043

Budnik, V. (1996). Synapse maturation and structural plasticity at Drosophila neuromuscular junctions. Current Opinion in Neurobiology, 6(6), 858–867. 10.1016/s0959-4388(96)80038-9

Camilleri-Robles, C., Amador, R., Tiebe, M., Teleman, A. A., Serras, F., Guigó, R., & Corominas, M. (2024). Long non-coding RNAs involved in Drosophila development and regeneration. NAR genomics and bioinformatics, 6(3), lqae091. 10.1093/nargab/lqae091

Cassé, C., Giannoni, F., Nguyen, V. T., Dubois, M. F., & Bensaude, O. (1999). The transcriptional inhibitors, actinomycin D and α-amanitin, activate the HIV-1 promoter and favor phosphorylation of the RNA polymerase II C-terminal domain. Journal of Biological Chemistry, 274(23), 16097–16106. 10.1074/jbc.274.23.16097

Castells-Nobau, A., Nijhof, B., Eidhof, I., Wolf, L., Scheffer-de Gooyert, J. M., Monedero, I., Torroja, L., van der Laak, J. A. W. M., & Schenck, A. (2017). Two Algorithms for High-throughput and Multi-parametric Quantification of Drosophila Neuromuscular Junction Morphology. Journal of Visualized Experiments : JoVE, (123), 55395. 10.3791/55395

Chang, N., Wells, J. N., Wang, A. Y., Schofield, P., Huang, Y., Truong, V. H., Simoes-Costa, M., & Feschotte, C. (2025). Gag proteins encoded by endogenous retroviruses are required for zebrafish development. Proceedings of the National Academy of Sciences, 122(18), e2411446122. 10.1073/pnas.2411446122

Chung, W. J., Okamura, K., Martin, R., & Lai, E. C. (2008). Endogenous RNA interference provides a somatic defense against Drosophila transposons. Current Biology, 18(11), 795– 802. 10.1016/j.cub.2008.05.006

Cui, M., Wang, Y., Xu, M., Wei, H., Bai, B., Chen, R., … & Li, M. (2026). Long Noncoding RNA SDRG Regulates Drosophila Neuromuscular Synapse Development by Modulating Frequenin 2 Through Coracle. The FASEB Journal, 40(12), e72024. 10.1096/fj.202601174R

Davie, K., Janssens, J., Koldere, D., De Waegeneer, M., Pech, U., Kreft, Ł., Aibar, S., Makhzami, S., Christiaens, V., Bravo González-Blas, C., Poovathingal, S., Hulselmans, G., Spanier, K. I., Moerman, T., Vanspauwen, B., Geurs, S., Voet, T., Lammertyn, J., Thienpont, B., … Aerts, S. (2018). A Single-Cell Transcriptome Atlas of the Aging Drosophila Brain. Cell, 174(4), 982–998.e20. 10.1016/j.cell.2018.05.057

Derrien, T., Johnson, R., Bussotti, G., Tanzer, A., Djebali, S., Tilgner, H., Guernec, G., Martin, D., Merkel, A., Knowles, D. G., Lagarde, J., Veeravalli, L., Ruan, X., Ruan, Y., Lassmann, T., Carninci, P., Brown, J. B., Lipovich, L., Gonzalez, J. M., … Guigó, R. (2012). The GENCODE v7 catalog of human long noncoding RNAs: Analysis of their gene structure, evolution, and expression. Genome Research, 22(9), 1775–1789. 10.1101/gr.132159.111

Di Franco, C., Pisano, C., Dimitri, P., Gigliotti, S., & Junakovic, N. (1989). Genomic distribution of copia-like transposable elements in somatic tissues and during development of Drosophila melanogaster. Chromosoma, 98(6), 402–410. 10.1007/BF00292785

Gu, W., Shirayama, M., Conte, D., Vasale, J., Batista, P. J., Claycomb, J. M., Moresco, J. J., Youngman, E. M., Keys, J., Stoltz, M. J., Chen, C. C. G., Chaves, D. A., Duan, S., Kasschau, K. D., Fahlgren, N., Yates, J. R., Mitani, S., Carrington, J. C., & Mello, C. C. (2009). Distinct Argonaute-Mediated 22G-RNA Pathways Direct Genome Surveillance in the C. elegans Germline. Molecular Cell, 36(2), 231–244. 10.1016/j.molcel.2009.09.020

Hayward, A., & Gilbert, C. (2022). Transposable elements. Current Biology, 32(17), R904–R909. 10.1016/j.cub.2022.07.044

Imamichi, T., Conrads, T. P., Zhou, M., Liu, Y., Adelsberger, J. W., Veenstra, T. D., & Lane, H. C. (2005). A transcription inhibitor, actinomycin D, enhances HIV-1 replication through an interleukin-6-dependent pathway. JAIDS Journal of Acquired Immune Deficiency Syndromes, 40(4), 388–397. 10.1097/01.qai.0000179466.25700.2f

Inagaki, S., Numata, K., Kondo, T., Tomita, M., Yasuda, K., Kanai, A., & Kageyama, Y. (2005). Identification and expression analysis of putative mRNA-like non-coding RNA in Drosophila. Genes to Cells, 10(12), 1163–1173. 10.1111/j.1365-2443.2005.00910.x

Klumpe, S., Senti, K. A., Beck, F., Sachweh, J., Hampoelz, B., Ronchi, P., Oorschot, V., Brandstetter, M., Yeroslaviz, A., Briggs, J. A. G., Brennecke, J., Beck, M., & Plitzko, J. M. (2025). In-cell structure and snapshots of copia retrotransposons in intact tissue by cryo-ET. Cell, 188(8), 2094–2110.e18. 10.1016/j.cell.2025.02.003

Koh, Y. H., Popova, E., Thomas, U., Griffith, L. C., & Budnik, V. (1999). Regulation of DLG localization at synapses by CaMKII-dependent phosphorylation. Cell, 98(3), 353–363. 10.1016/s0092-8674(00)81964-9

Lander, E. S., Linton, L. M., Birren, B., Nusbaum, C., Zody, M. C., Baldwin, J., Devon, K., Dewar, K., Doyle, M., FitzHugh, W., Funke, R., Gage, D., Harris, K., Heaford, A., Howland, J., Kann, L., Lehoczky, J., LeVine, R., McEwan, P., McKernan, K., … International Human Genome Sequencing Consortium (2001). Initial sequencing and analysis of the human genome. Nature, 409(6822), 860–921. 10.1038/35057062

Latzel, V., Puy, J., Thieme, M., Bucher, E., Götzenberger, L., & de Bello, F. (2023). Phenotypic diversity influenced by a transposable element increases productivity and resistance to competitors in plant populations. Journal of Ecology, 111(11), 2376–2387. 10.1111/1365-2745.14185

Lefebvre, F. A., Benoit Bouvrette, L. P., Perras, L., Blanchet-Cohen, A., Garnier, D., Rak, J., & Lécuyer, É. (2016). Comparative transcriptomic analysis of human and Drosophila extracellular vesicles. Scientific Reports, 6(1), 27680. 10.1038/srep27680

Li, W., Prazak, L., Chatterjee, N., Grüninger, S., Krug, L., Theodorou, D., & Dubnau, J. (2013). Activation of transposable elements during aging and neuronal decline in Drosophila. Nature Neuroscience, 16(5), 529–531. 10.1038/nn.3368

Lo Piccolo, L., & Yamaguchi, M. (2017). RNAi of arcRNA hsrω affects sub-cellular localization of Drosophila FUS to drive neurodiseases. Experimental Neurology, 292, 125–134. 10.1016/j.expneurol.2017.03.011

Lu, T. C., Brbić, M., Park, Y. J., Jackson, T., Chen, J., Kolluru, S. S., Qi, Y., Katheder, N. S., Cai, X. T., Lee, S., Chen, Y. C., Auld, N., Liang, C. Y., Ding, S. H., Welsch, D., D’Souza, S., Pisco, A. O., Jones, R. C., Leskovec, J., … Li, H. (2023). Aging Fly Cell Atlas identifies exhaustive aging features at cellular resolution. Science, 380(6650), eadg0934. 10.1126/science.adg0934

M’Angale, P. G., Lemieux, A., Liu, Y., Wang, S., Zinter, M., Alegre, G., … & Thomson, T. (2025). Capsid transfer of the retrotransposon Copia controls structural synaptic plasticity in Drosophila. PLoS Biology, 23(2), e3002983. 10.1371/journal.pbio.3002983

Marchese, F. P., Raimondi, I., & Huarte, M. (2017). The multidimensional mechanisms of long noncoding RNA function. Genome Biology, 18(1), 206. 10.1186/s13059-017-1348-2

Mouse Genome Sequencing Consortium, Waterston, R. H., Lindblad-Toh, K., Birney, E., Rogers, J., Abril, J. F., Agarwal, P., Agarwala, R., Ainscough, R., Alexandersson, M., An, P., Antonarakis, S. E., Attwood, J., Baertsch, R., Bailey, J., Barlow, K., Beck, S., Berry, E., Birren, B., Bloom, T., … Lander, E. S. (2002). Initial sequencing and comparative analysis of the mouse genome. Nature, 420(6915), 520–562. 10.1038/nature01262

Niu, X. M., Xu, Y. C., Li, Z. W., Bian, Y. T., Hou, X. H., Chen, J. F., Zou, Y. P., Jiang, J., Wu, Q., Ge, S., Balasubramanian, S., & Guo, Y. L. (2019). Transposable elements drive rapid phenotypic variation in Capsella rubella. Proceedings of the National Academy of Sciences of the United States of America, 116(14), 6908–6913. 10.1073/pnas.1811498116

Nojima, T., & Proudfoot, N. J. (2022). Mechanisms of lncRNA biogenesis as revealed by nascent transcriptomics. Nature Reviews Molecular Cell Biology, 23(6), 389–406. 10.1038/s41580-021-00447-6

Pasyukova, E. G., & Nuzhdin, S. V. (1993). Doc and copia instability in an isogenic Drosophila melanogaster stock. Molecular and General Genetics MGG, 240(2), 302–306. 10.1007/BF00277071

Pasyukova, E., Nuzhdin, S., Li, W., & Flavell, A. J. (1997). Germ line transposition of the copia retrotransposon in Drosophila melanogaster is restricted to males by tissue-specific control of copia RNA levels. Molecular and General Genetics MGG, 255(1), 115–124. 10.1007/s004380050479

Percharde, M., Lin, C. J., Yin, Y., Guan, J., Peixoto, G. A., Bulut-Karslioglu, A., Biechele, S., Huang, B., Shen, X., & Ramalho-Santos, M. (2018). A LINE1-Nucleolin Partnership Regulates Early Development and ESC Identity. Cell, 174(2), 391–405.e19. 10.1016/j.cell.2018.05.043

Perkins, L. A., Holderbaum, L., Tao, R., Hu, Y., Sopko, R., McCall, K., Yang-Zhou, D., Flockhart, I., Binari, R., Shim, H. S., Miller, A., Housden, A., Foos, M., Randkelv, S., Kelley, C., Namgyal, P., Villalta, C., Liu, L. P., Jiang, X., Huan-Huan, Q., … Perrimon, N. (2015). The Transgenic RNAi Project at Harvard Medical School: Resources and Validation. Genetics, 201(3), 843–852. 10.1534/genetics.115.180208

Port, F., Chen, H. M., Lee, T., & Bullock, S. L. (2014). Optimized CRISPR/Cas tools for efficient germline and somatic genome engineering in Drosophila. Proceedings of the National Academy of Sciences of the United States of America, 111(29). 10.1073/pnas.1405500111

Silva, G. M., Kaplan, A. L., Park, C. R., Picone, J. A., Kim, R. K., Truby, N. L., … & Hamilton, P. J. (2026). Transposable elements are dynamically regulated in medium spiny neurons and may contribute to the molecular and behavioral adaptations to cocaine. Biological Psychiatry, 100(3), 270–281. 10.1016/j.biopsych.2025.07.014

Siudeja, K., van den Beek, M., Riddiford, N., Boumard, B., Wurmser, A., Stefanutti, M., Lameiras, S., & Bardin, A. J. (2021). Unraveling the features of somatic transposition in the Drosophila intestine. The EMBO Journal, 40(9), e106388. 10.15252/embj.2020106388

Stein, P., Rozhkov, N. V., Li, F., Cárdenas, F. L., Davydenk, O., Vandivier, L. E., Gregory, B. D., Hannon, G. J., & Schultz, R. M. (2015). Essential Role for Endogenous siRNAs during Meiosis in Mouse Oocytes. PLoS Genetics, 11(2), e1005013. 10.1371/journal.pgen.1005013

Treiber, C. D., & Waddell, S. (2017). Resolving the prevalence of somatic transposition in *Drosophila*. eLife, 6, e28297. 10.7554/eLife.28297

Wang, W., Han, B. W., Tipping, C., Ge, D. T., Zhang, Z., Weng, Z., & Zamore, P. D. (2015). Slicing and Binding by Ago3 or Aub Trigger Piwi-Bound piRNA Production by Distinct Mechanisms. Molecular Cell, 59(5), 819–830. 10.1016/j.molcel.2015.08.007

Xiao, C., M’Angale, P. G., Wang, S., Lemieux, A., & Thomson, T. (2023). Identifying new players in structural synaptic plasticity through dArc1 interrogation. iScience, 26(11), 108048. 10.1016/j.isci.2023.108048

Xue, Z., Ren, M., Wu, M., Dai, J., Rong, Y. S., & Gao, G. (2014). Efficient gene knock-out and knock-in with transgenic Cas9 in Drosophila. G3: Genes, Genomes, Genetics, 4(5), 925–929. 10.1534/g3.114.010496

Zhang, X., Jiang, Q., Li, J., Zhang, S., Cao, Y., Xia, X., Cai, D., Tan, J., Chen, J., & Han, J. D. J. (2022). KCNQ1OT1 promotes genome-wide transposon repression by guiding RNA–DNA triplexes and HP1 binding. Nature Cell Biology, 24(11), 1617–1629. 10.1038/s41556-022-01008-5

